# Differential composition of lymphocyte subpopulations and activation between the hypertensive Bph/2 and normotensive Bpn/3 mouse strains

**DOI:** 10.1101/2024.10.01.616072

**Authors:** Devon Dattmore, Paiton McDonald, Afrin Chowdhury, Saamera Awali, Allison P. Boss, Yining Jin, Lisa Sather, D. Adam Lauver, Cheryl E. Rockwell

**Affiliations:** Department of Pharmacology and Toxicology; Department of Animal Science; Integrated Pharmacological Sciences Training Program; Department of Food Science and Human Nutrition; Applied Immunology Center for Education and Research Michigan State University, East Lansing, MI 48824

## Abstract

Numerous studies point to a role for the immune system in various animal models of hypertension. However, little is known about the immune system of Bph/2 mice, a spontaneously hypertensive strain. To address this, we conducted a comprehensive comparison of immune cell composition and response to polyclonal T cell activation in hypertensive Bph/2 mice and normotensive Bpn/3 control mice. We quantified immune cell populations by flow cytometry from spleen and inguinal, brachial and mesenteric lymph nodes. While composition of myeloid immune cell types was largely comparable between strains, we observed differences in B and T cell subpopulations. Specifically, we found an increased percentage of IgM+ IgD^Lo^ and IgM+ IgD-B cells in Bph/2 mice, suggesting greater baseline B cell activation. In addition, we observed a decreased percentage of CD4 effector memory T cells and CD8 central memory T cells. The diminished proportion of memory T cells in Bph/2 mice correlated with decreased proliferation and cytokine response of splenic T cells to polyclonal T cell activation. In splenic T cells from Bph/2 mice 24 h after activation we observed a pronounced decrease in the majority of T cell cytokines. At 120 h after activation, the Th1 and Th17 cytokine responses of splenic T cells from Bph/2 mice were decreased, but other T cell cytokines were largely comparable. Overall, the data suggest a decreased percentage of memory T cells in Bph/2 mice that correlates with markedly diminished proliferation and a reduced cytokine response to polyclonal activation.

## Introduction

The Bph/2 mouse strain was originally generated by Dr. Gunther Schlager with the purpose of creating a genetic model of hypertension in mice. To achieve this, Dr. Schlager crossed eight different inbred strains of mice, including LP/J, SJL/J, BALB/cJ, C57BL/6J, 129/J, CBA/J, RF/J and BDP/J [1; 2]. After creating the 8-way cross, he bred the resulting progeny together for three generations before stratifying them based on blood pressure. Using this approach, Dr. Schlager created numerous mouse strains— many of which continue to be used. These include Bpl/1 mice (which have low blood pressure), Bph/2 (which have high blood pressure) and Bpn/3 (which are a normotensive control strain). The differences in blood pressure among these strains occurs early in life and is substantial. The systolic blood pressures of Bph/2 mice is more than 30 mm Hg greater than those of Bpn/3 mice [1]. The difference increased to approximately 50 mm Hg when comparing Bph/2 and Bpl/1 mice. The elevated blood pressure in Bph/2 mice was found to occur in mice as young as 6 weeks of age. Schlager’s group also found a slightly elevated heart rate in Bph/2 mice as compared to Bpn/3 mice [1].

The development of hypertension is notably complex with roles for the sympathetic nervous system, vascular contraction/stiffness and kidney function, among other factors. In addition to these, the immune system is emerging as another possible element that may contribute to blood pressure regulation. Interest in the potential role of the immune system on blood pressure stems back many decades to studies conducted in the 1960’s that showed that immunosuppressive drugs were protective in certain animal models of hypertension and that immune cells from hypertensive animals can transfer hypertension into non-hypertensive animals [3; 4]. More recent studies have shown that angiotensin II-driven hypertension is diminished in Rag 1 knockout mice (Rag1 is a gene necessary for T cell development [5]). Likewise, the DOCA salt model of hypertension is also less pronounced in Rag1-/-mice. There have been analogous studies performed in rats demonstrating that high salt-driven hypertension in Dahl S rats is decreased in the absence of Rag1 or CD247, a gene necessary for a functional T cell receptor [6; 7].

While these data point to a role for the immune system in at least some models of hypertension, the immune system in Bph/2 mice has not been well characterized. Thus, the purpose of the present study was to address this. Toward this end, we collected leukocytes from several different lymphoid compartments, including spleen and inguinal, brachial and mesenteric lymph nodes. We compared immune cell composition between the Bph/2 (hypertensive) and Bpn/3 (normotensive) strains. We also investigated potential differences in immune cell activation in splenocytes collected between the two different strains.

## Methods

### Animals

Male Bph/2 and Bpn/3 mice were purchased from Jackson Laboratories (Bar Harbor, ME). The animals were randomized into cages prior to a 1-week acclimatization period and were housed in specific pathogen free conditions. The mice were provided food and water ad libitum and maintained on a 12 h light-dark cycle. Tissues were collected from the mice at 12 weeks of age. This study protocol was approved by the Institutional Animal Care and Use Committee (IACUC) at Michigan State University and conducted in accordance with the NIH Guide for the Care and Use of Animals.

### Tail-Cuff Plethysmography

Blood pressure was measured in conscious mice by tail-cuff plethysmography using a RTBP1001 tail-cuff blood pressure system (CODA-6, Kent Scientific, Torrington, CT). All mice were acclimatized to both handling and the blood pressure measurement system ahead of the experimental measurements. Of the 25 cycles measured, the first 15 were used as acclimation, and the remaining 10 were averaged to provide the final data point for each animal.

### Tissue Collection

Spleen and inguinal, brachial and mesenteric lymph node tissues were aseptically collected from 12-week-old Bpn/3 or Bph/2 mice and processed into single-cell suspensions. The cells from inguinal, brachial and mesenteric lymph nodes were immediately analyzed via flow cytometry. Splenocytes were divided into two fractions: one that was immediately analyzed via flow cytometry and another that was cultured for immune cell activation and subsequent analysis.

### Activation of splenocytes

Splenocytes were activated with anti-CD3/anti-CD28/crosslinker (1.5 μg/mL anti-CD3, μg/mL anti-CD28, 1.5 μg/mL crosslinker) and cultured at 37°C, 5% CO_2_ in RPMI 1640 containing 100 U penicillin/mL, 100 U streptomycin/mL, 25 mM HEPES, 10 mM nonessential amino acids, 1 mM sodium pyruvate, and 10% fetal bovine serum (Biowest LLC, Kansas City, MO). The cell supernatants were collected 24 h and 120 h after activation and frozen at -80°C for cytokine analysis. Cells were collected 24 h after activation for quantification of activation markers by flow cytometry.

### Flow cytometry

Cells from spleen and inguinal, brachial and mesenteric lymph nodes were labeled using three different anti-body staining panels (a broad immune panel, a T cell-specific panel, and a B cell-specific panel). Splenic T cells activated with anti-CD3/anti-CD28 were analyzed using the T cell-specific panel (24-hours post activation). Cells were stained with Zombie Aqua Fixable Dye and labeled as described previously [8]. The following antibodies were used:

**Broad Immune Panel**
| Antigen | Fluorochrome | Dilution | Manufacturer |
| --- | --- | --- | --- |
| CD3 | AF488 | 1:100 | Biolegend |
| CD4 | APC/Cy7 | 1:100 | Biolegend |
| CD8 | PE/Cy5.5 | 1:100 | Invitrogen |
| CD11b | BV605 | 1:100 | Biolegend |
| CD11c | AF647 | 1:100 | Biolegend |
| CD19 | PE/Dazzle594 | 1:100 | Biolegend |
| CD45 | PerCP/Cy5.5 | 1:100 | Biolegend |
| F4/80 | AF700 | 1:100 | Biolegend |
| Gr-1 | PacBlue | 1:100 | Biolegend |
| I-Ab | PE/Cy7 | 1:100 | Biolegend |
| NK1.1 | BV711 | 1:100 | Biolegend |
| SiglecF | PE | 1:100 | Biolegend |

**B Cell Panel**
| Antigen | Fluorochrome | Dilution | Manufacturer |
| --- | --- | --- | --- |
| CD19 | PE/Dazzle594 | 1:100 | Biolegend |
| CD23 | PacBlue | 1:100 | Biolegend |
| CD69 | PE/Cy7 | 1:100 | Biolegend |
| CD86 | APC/Cy7 | 1:100 | Biolegend |
| IgD | BV605 | 1:100 | Biolegend |
| IgM | AF488 | 1:100 | Biolegend |
| CD138 | PE | 1:100 | Biolegend |

**T Cell Panel**
| Antigen | Fluorochrome | Dilution | Manufacturer |
| --- | --- | --- | --- |
| CD3 | AF488 | 1:100 | Biolegend |
| CD4 | APC/Cy7 | 1:100 | Biolegend |
| CD8 | PE/Cy5.5 | 1:100 | Invitrogen |
| CD25 | BV711 | 1:100 | Biolegend |
| CD44 | PE/Dazzle594 | 1:100 | Biolegend |
| CD62L | PacBlue | 1:100 | Biolegend |
| CD69 | AF700 | 1:100 | Biolegend |
| CD134 | BV605 | 1:100 | Biolegend |
| CD137 | PE/Cy7 | 1:100 | Biolegend |
| CD154 | APC | 1:100 | Biolegend |

## Broad Immune Panel

### Cell Proliferation

Splenocytes (25,000 cells/well) in a 96-well plate were labeled with IncuCyte Nuclight Rapid Red Cell Labeling kit following the manufacturer’s protocol (Sartorius, Gottingen, Germany). Fluorescence was quantified every 8 h using the Incucyte S3 Live-Cell Analysis System (Sartorius, Gottingen, Germany).

### Cytokine Analysis

Supernatants were shipped to Eve Technologies (Calgary, AB) for cytokine analysis. Each sample was analyzed by multiplex bead array using the MD32 and MD12-TH17 kits.

### Statistical Analysis

The mean ± SEM was determined for each treatment group in each experiment. For analyses with two experimental groups, a Student’s t test was used (Graphpad Prism 10). For analyses with more than two experimental groups and normally distributed data, a one-way or a two-way ANOVA was used (Graphpad Prism 10). For analyses with that were not normally distributed with multiple groups, the data were first log transformed prior to analysis by one-way or two-way ANOVA and a Dunnet’s post hoc test using Sigmaplot v12 (Systat Software, San Jose CA).

## Results

### Lower body weight, but increased blood pressure, in Bph/2 mice

We quantified blood pressure by tail cuff measurement. We opted for tail cuff measurements in this study instead of telemetery due to our concern regarding the impact of telemeter implants on baseline inflammation. As expected, Bph/2 mice had significantly elevated blood pressures as compared to the normotensive Bpn/3 control strain (Fig. 1). In contrast, heart rates were not significantly different between the two strains at this time point. We also observed a difference in body weight in which the Bpn/3 mice had significantly greater body mass as compared to Bph/2 mice. Likewise, we observed a proportional difference in spleen weights such that the spleen-to-body weight ratio was equal between the two strains.

**Figure 1.**
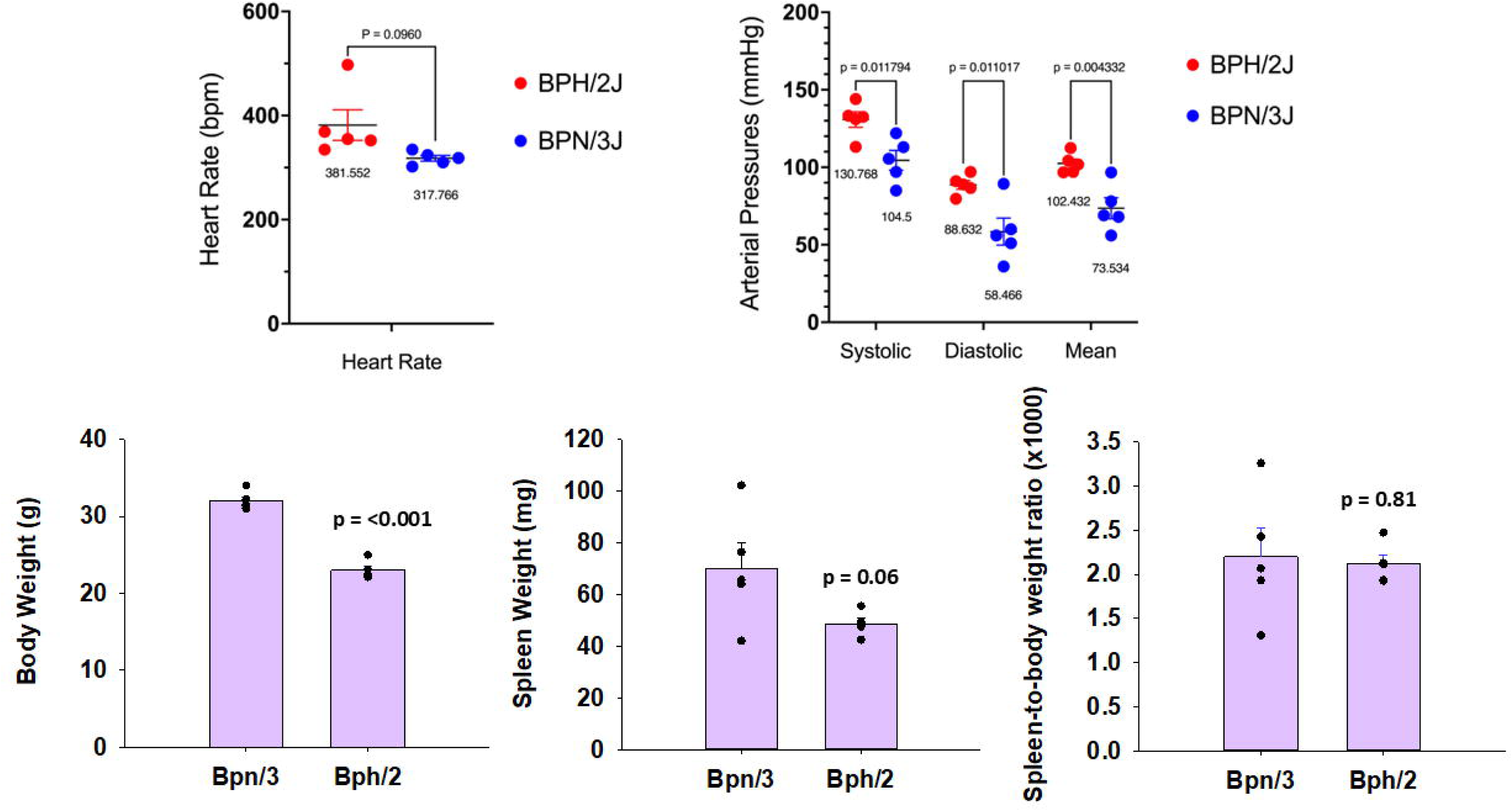
Bph/2 mice have elevated arterial blood pressure and lower body weights. Blood pressure was determined by tail cuff plethysmography. Shown in the graphs are the heart rates, systolic, diastolic, and mean arterial pressures, body weights, spleen weights, and body-to-spleen weight ratios for Bpn/3 and Bph/2 mice. Data are presented as the mean ± SEM, n = 5. * p<0.05 and ***p<0.001 as determined by unpaired Student’s t-test.

### With the exception of neutrophils, composition of myeloid cell populations are largely comparable between Bph/2 and Bpn/3 mice

We assessed the composition of various immune cell populations across multiple immune compartments, including spleen as well as inguinal, brachial and mesenteric lymph nodes. Of the granulocyte populations, the percentage of eosinophils was fairly comparable between Bpn/3 and Bph/2 mice in the different compartments (Fig. 2). In contrast, the percentage of neutrophils in the spleens of Bph/2 mice was significantly greater than those from Bpn/3 mice. It should be noted that we were unable to quantify neutrophils in any of the lymph nodes due to low numbers. With respect to the antigen presenting cells, there was a modestly lower percentage of dendritic cells in mesenteric lymph nodes of Bph/2 mice as compared to Bpn/3. Likewise, we also observed a decreased percentage of macrophages in mesenteric lymph nodes of Bph/2 mice. However, there were no differences in the proportion of dendritic cell or macrophage populations in spleen, inguinal or brachial lymph nodes. Taken together, the data indicate that Bph/2 mice have more than double the percentage of splenic neutrophils compared to Bpn/3 mice, whereas most of the other myeloid populations were comparable between Bph/2 and Bpn/3 mice.

**Figure 2.**
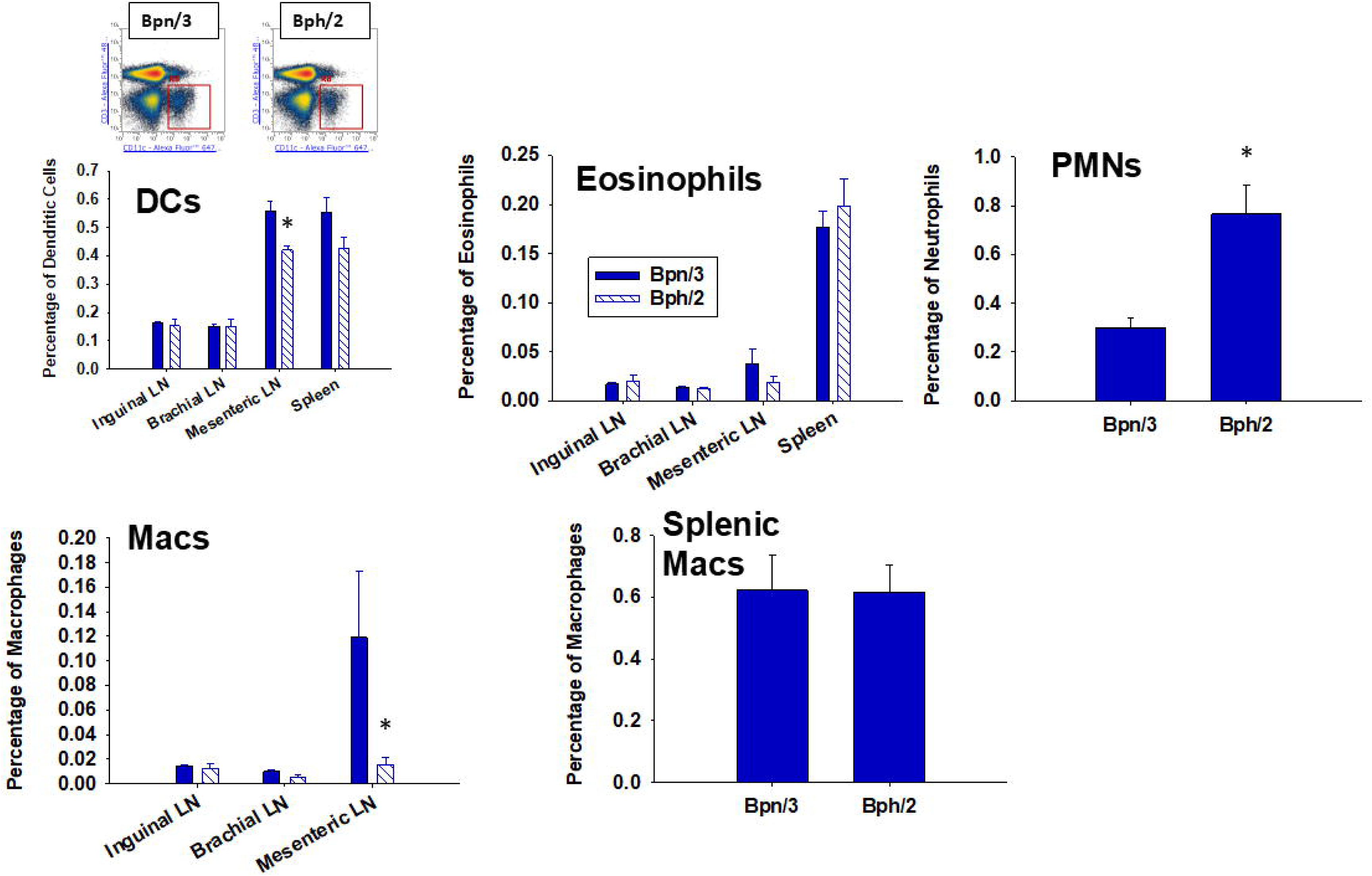
Bpn/2 mice have elevated percentages of splenic neutrophils and decreased percentages of macrophages and dendritic cells in the mesenteric lymph nodes. Splenocytes and lymph nodes (brachial, inguinal, and mesenteric) were isolated from twelve-week-old Bph/2 and Bpn/3 mice, labeled with fluorochrome-conjugated antibodies (Table 1) and analyzed by flow cytometry. Representative flow density plots are shown above, and graphs shown below illustrate the mean ± SEM values for each respective group, n = 5. The graphs depict dendritic cells, eosinophils, neutrophils (splenic only), macrophages, and splenic macrophages as percentages of the live cell population. * p<0.05 as determined by unpaired Student’s t-test.

### Bpn/3 mice have a decreased CD4/CD8 ratio in inguinal and brachial lymph nodes

In wild-type C57Bl/6 mice (which is the most widely used mouse strain in immunology studies), the ratio of CD4 to CD8 T cells tends to remain fairly stable with a higher percentage of CD4 T cells relative to CD8 T cells. Consistent with this, we observed a greater proportion of CD4 T cells compared to CD8 T cells in every lymphoid compartment we investigated in Bph/2 mice (Fig. 3). The proportion of CD4 T cells was also greater than that of CD8 T cells in the spleens and mesenteric lymph nodes of Bpn/3 mice. In contrast, the percentages of CD4 T cells and CD8 T cells were largely equivalent in the inguinal and brachial lymph nodes of Bpn/3 mice. Overall, we found a lower CD4 to CD8 T cell ratio in the spleen and mesenteric lymph nodes of Bpn/3 mice, which suggests a possible difference in T cell development between the two strains.

**Figure 3.**
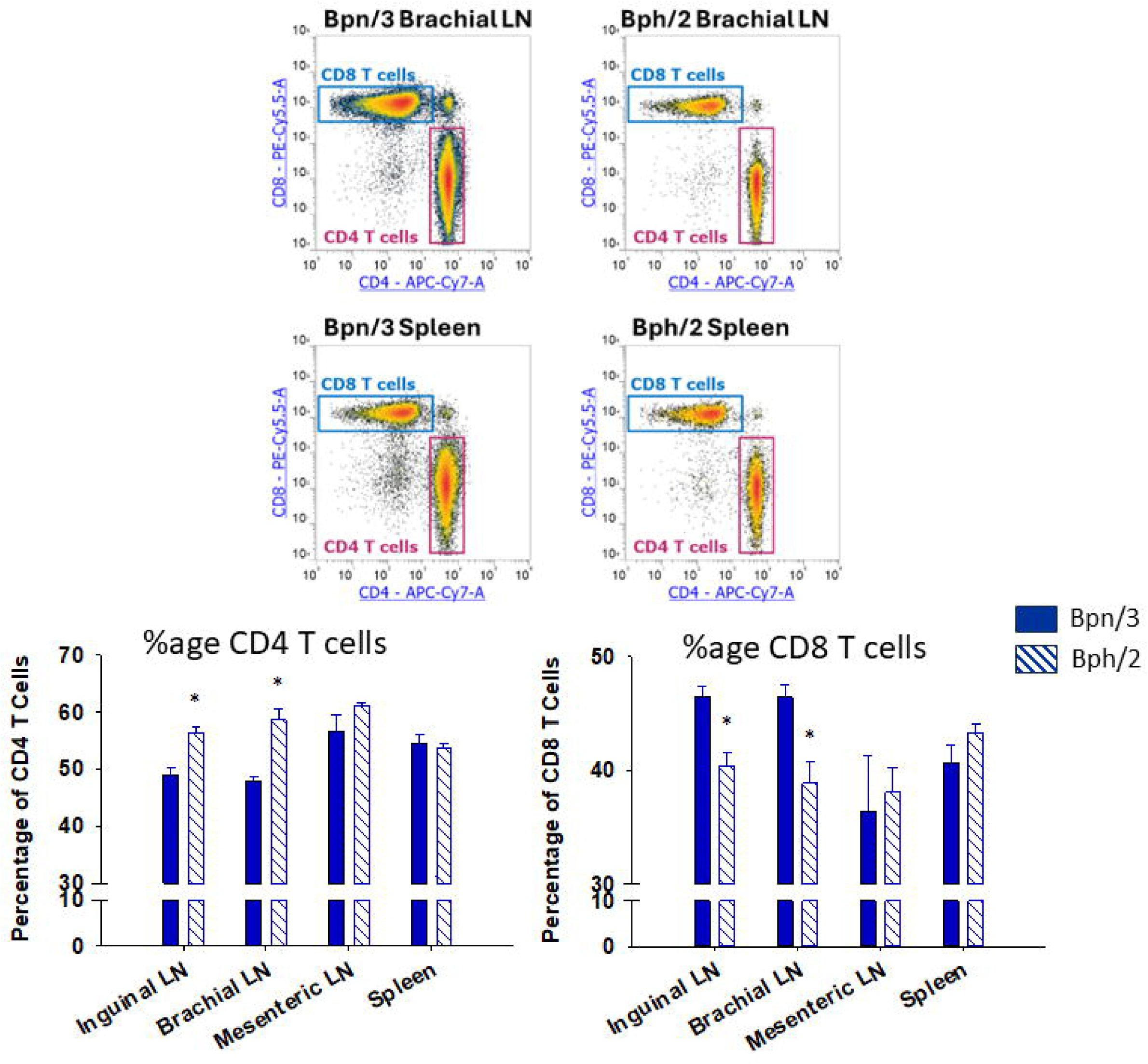
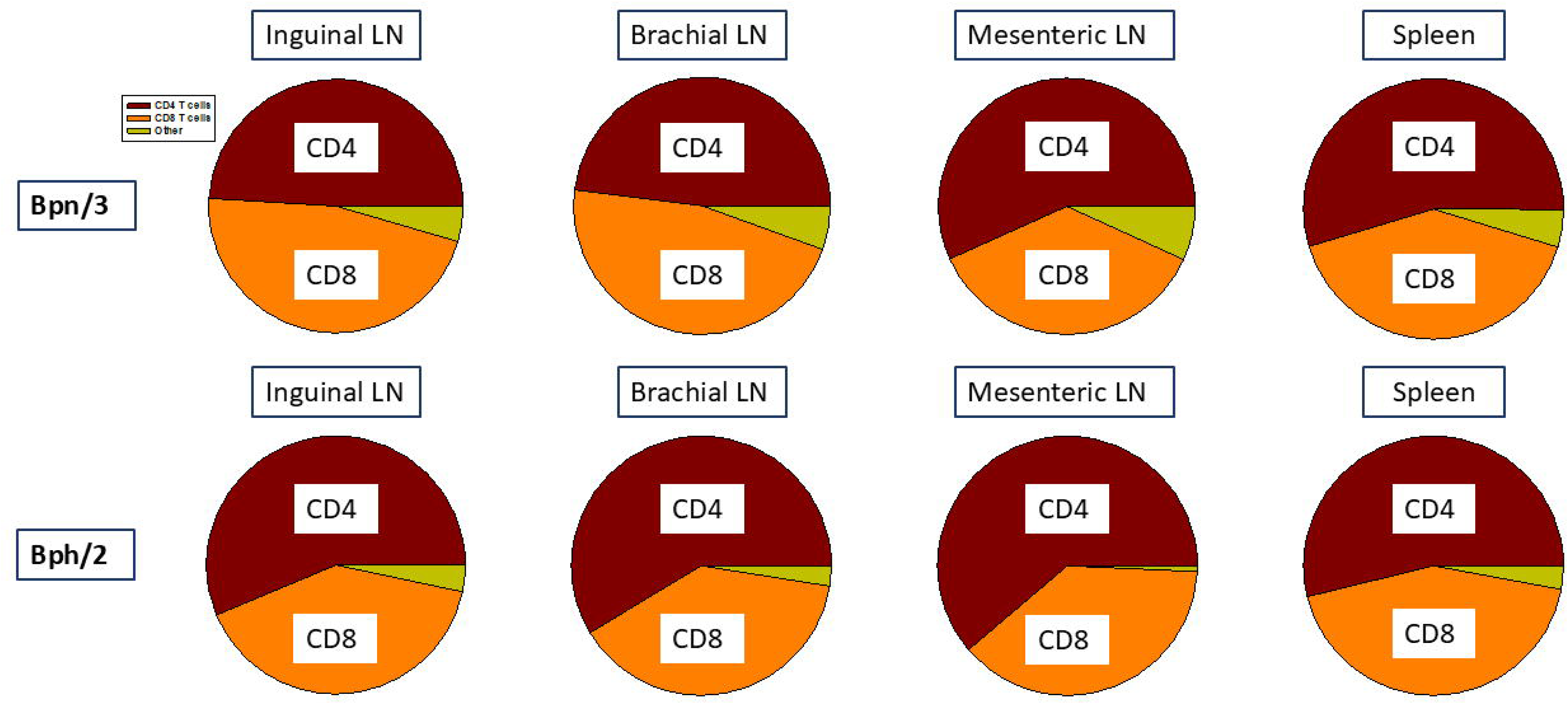
Bph/2 mice have altered percentages of CD4^+^ and CD8^+^ T cells in brachial and inguinal lymph nodes compared to Bpn/3 mice. Single-cell suspensions were extracted from spleens and lymph nodes of twelve-week-old Bpn/3 and Bph/2 mice, labeled with fluorochrome-conjugated antibodies (Table 1) and analyzed by flow cytometry. (A) CD4^+^ and CD8^+^ T cell percentages relative to CD3^+^ T cell population. Representative flow density plots are shown above, and graphs shown below illustrate the mean ± SEM values for each respective group, n = 5. (B) Pie graphs showing the relative proportions of CD4^+^ and CD8^+^ T cells in relation to the whole T cell population. Other refers to T cell populations that were CD4+ CD8+ double-positive or CD4-CD8-double-negative *p<0.05, **p<0.01, ****p<0.0001 as determined by unpaired Student’s t-test.

### Bph/2 mouse spleens have a lower percentage of IgM+ IgD^Hi^, but a greater percentage of IgM+ IgD^Lo^ and IgM+ IgD-B cells

One of the primary functions of B cells is to express antibodies, such as IgD and IgM. Newly matured B cells express high levels of IgD and low levels of IgM. When B cells become activated, the ratio of IgM to IgD expression shifts. Activated B cells undergo class switching to solely express IgM with a transition period between where B cells express high levels of IgM and low levels of IgD. We observed notable differences in antibody expression in B cells in the spleens of Bph/2 vs. Bpn/3 mice. Specifically, we found Bph/2 spleens showed a lower percentage of IgM+ IgD^Hi^ B cells with a concurrent increased percentage of IgM+ IgD^Lo^ and IgM+ IgD-B cells. (Fig. 4) In contrast, the percentage of IgM-IgD-B cells was roughly comparable in spleen, but was lower in the brachial lymph nodes of Bph/2 mice. The percentage of IgM+ IgD^Lo^ CD23-B cells, which would include marginal zone B cells was equivalent between the two strains in all lymphoid compartments. Likewise, expression of the B cell activation markers CD86 and CD69 (expressed by recently-activated B cells) was largely equivalent between strains with the exception of a low percentage of CD86+ B cells in the mesenteric lymph nodes of Bph/2 mice. B cells also have the ability to differentiate into plasma cells, a cell type that produces large quantities of antibody. Although we included included a marker for plasma cells in our panel (CD138), the percentage of plasma cells was too low to reliably measure. Collectively, these data suggest that splenic B cells from Bph/2 mice are more “developed” and have undergone greater class switching as compared to those from Bpn/3 mice.

**Figure 4.**
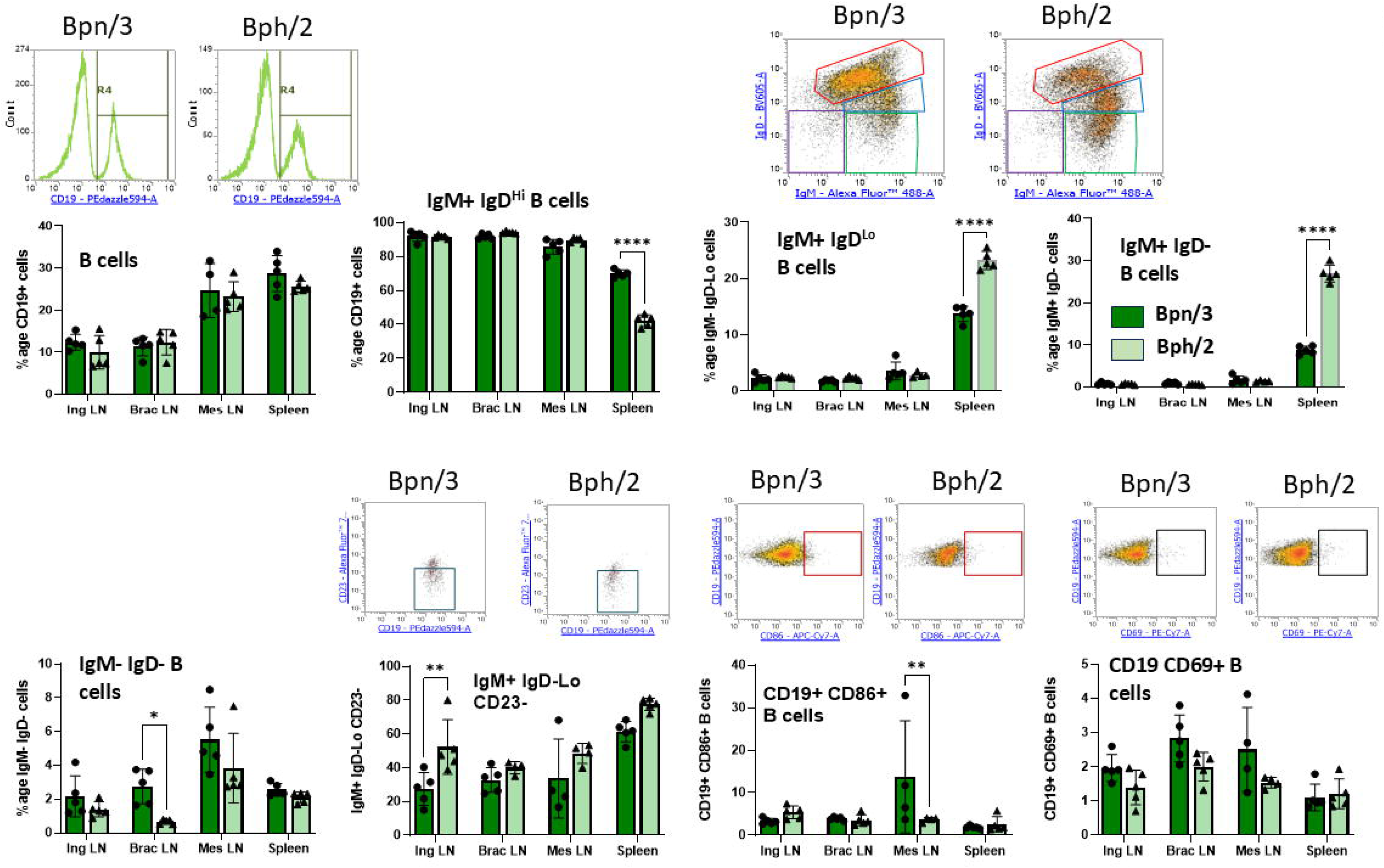
Lower percentage of IgM+ IgD-Lo and IgM+ IgD-B cells in spleens of Bph/2 mice. Single-cell suspensions were extracted from spleens and lymph nodes of twelve-week-old Bpn/3 and Bph/2 mice, labeled with fluorochrome-conjugated antibodies (Table 1) and analyzed by flow cytometry. The following B cell populations were quantified: the pan B cell population (CD19^+^), IgM^+^ IgD^Hi^ B cells (red gates), IgM^+^ IgD^Lo^ B cells (blue gates), IgM^+^ IgD-B cells (green gates), and IgM-IgD-B cells (purple gates) B cells, IgM^+^ IgD^Lo^ CD23-B cells (which includes marginal zone B cells) and recently activated (CD86^+^ and CD69^+^) B cells. Representative flow density plots are shown above, and graphs shown below illustrate the mean ± SEM values for each respective group, n = 5. *p<0.05 **p<0.01 and ****p<0.0001 as determined by unpaired Student’s t-test.

### Bph/2 mice have a pronounced decrease in the effector memory CD4 T cell population as compared to Bpn/3 mice

In addition to investigating effects on CD4 and CD8 T cell ratios, we also quantified T cell memory cell populations. The two dominant memory cell populations in lymphoid organs are effector memory T cells, which have a fast, robust cytokine response to stimulation, and central memory T cells, which have an intermediate response between effector memory T cells and naïve T cells. Naïve T cells have a slow, modest response to activation in comparison to memory cells. We found a substantially lower percentage of CD4 effector memory T cells in the Bph/2 mice as compared to Bpn/3 mice with a statistically significant difference in all lymphoid organs examined (Fig. 5a). In contrast, we did not observe any difference in the percentages of effector memory T cells within the CD8 T cell population, but rather, we observed a significantly lower percentage of central memory CD8 T cells that was consistent across all compartments (Fig. 5b). Taken together, the data indicate a reduction in the proportion of memory T cell populations in Bph/2 mice.

**Figure 5.**
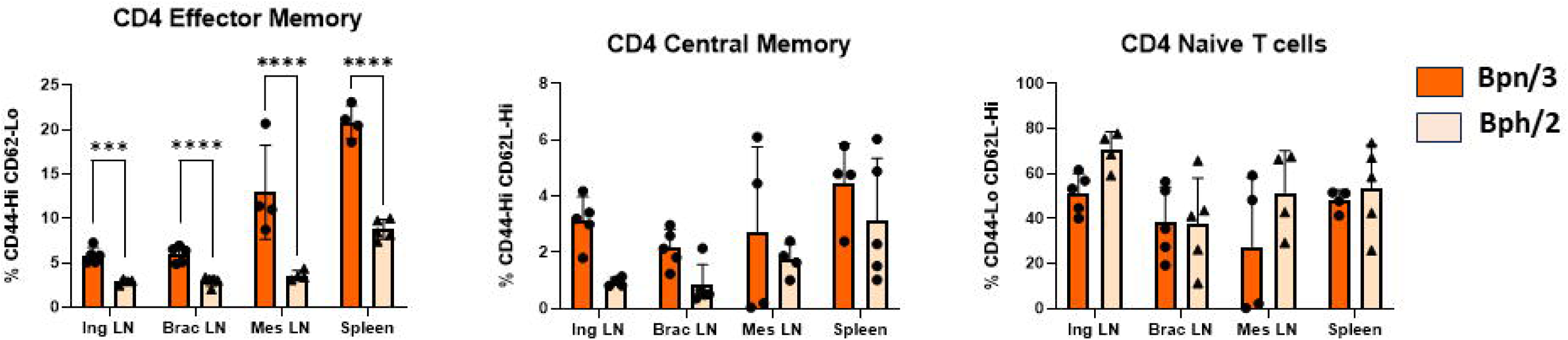
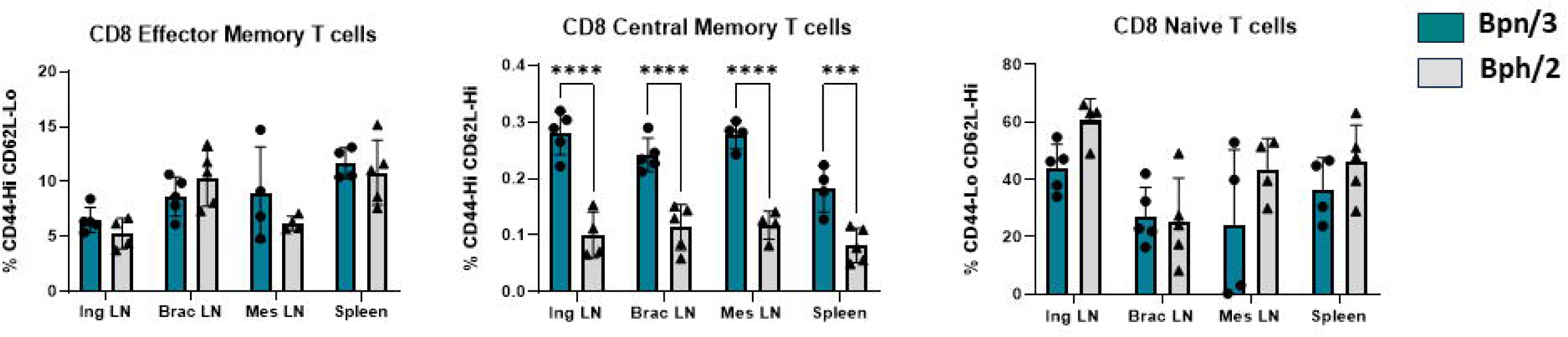
Lower percentages of CD4 effector memory T cells and CD8 central memory T cells in Bph/2 mice. Single-cell suspensions were extracted from spleens and lymph nodes of twelve-week-old Bpn/3 and Bph/2 mice, labeled with fluorochrome-conjugated antibodies (Table 1) and analyzed by flow cytometry. Naïve (CD62L^Hi^ CD44^Lo^), effector memory (CD62L^Lo^ CD44^Hi^)and central memory (CD62L^Hi^ CD44^Hi^) T cell subsets were quantified in the A) CD4 and B) CD8 T cell populations. No differences were observed in the effector memory (CD44^Hi^ CD62L^Lo^) or naïve (CD44^Lo^ CD62L^Hi^) CD8^+^ T cells. Bars are presented as the mean ± SEM. n = 5. ***p<0.001 and ****p<0.0001 as determined by unpaired Student’s t-test.

### Diminished proliferation in splenic T cells from Bph/2 mice activated with a polyclonal activator

In addition to assessing baseline homeostatic differences in immune cell composition between Bph/2 and Bpn/3 mice, we also compared the impact of polyclonal T cell activation (anti-CD3/anti-CD28) on splenic T cells from the two strains. For these studies, we used a freshly isolated splenocyte preparation activated with anti-CD3/anti-CD28. T cell activation results in a number of functional changes, including induction of proliferation, upregulation of certain cell surface receptors and an enormous increase in cytokine secretion. Notably, we observed markedly decreased proliferation in the cells derived from Bph/2 mice as compared to those from Bpn/3 mice (Fig. 6).

**Figure 6.**
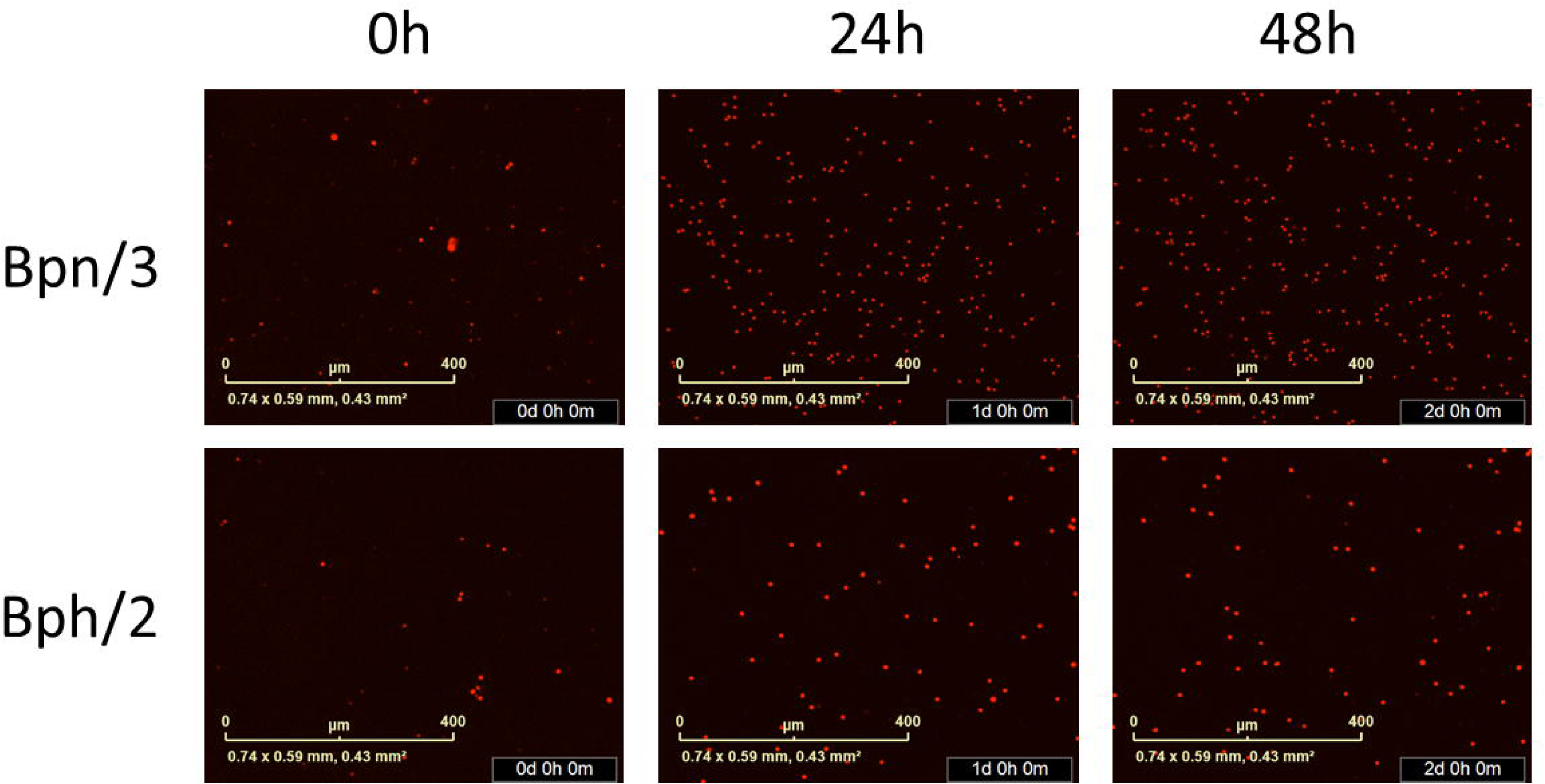
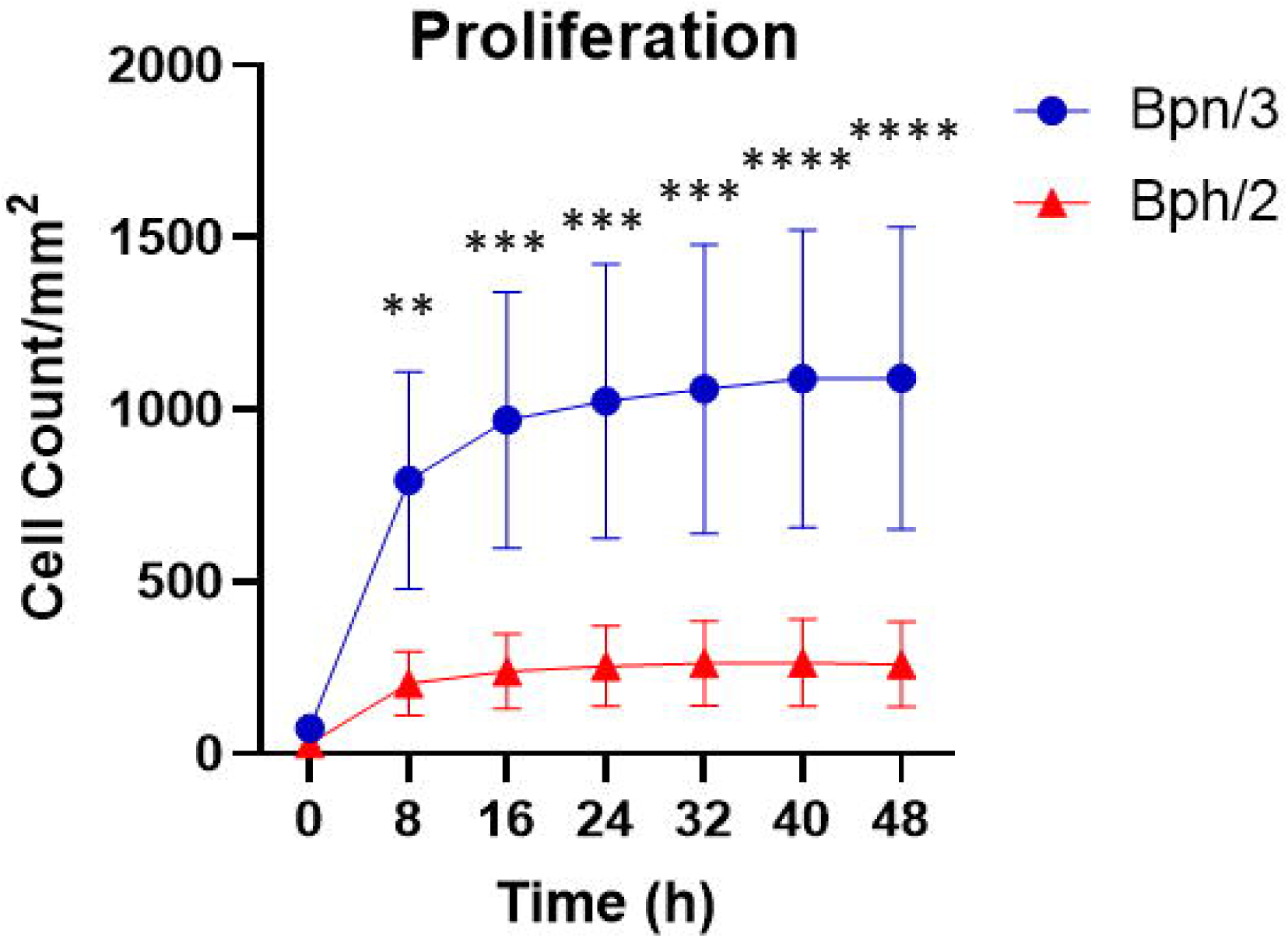
Decreased proliferation by activated splenic T cells from Bph/2 mice. Freshly isolated splenocytes were cultured (2.5×10^4^ cells/well), stained with IncuCyte Nuclight Rapid Red reagent, and cultured for 48 h. Fluorescence was quantified every 8 h using the Incucyte S3 Live-Cell Analysis System. A) Representative images of cells from Bph/2 and Bpn/2 mice. B) Line graph depicting cell counts at 0 – 48 h post activation. The data are presented as the mean ± SEM, n=5. * p<0.05 as determined by 2-way ANOVA followed by Dunnett’s post hoc analysis. **p<0.01, ***p<0.001 and ****p<0.0001.

### Decreased upregulation of activation markers in CD8 T cells from Bph/2 mice

Concurrent with an increase in proliferation, T cell activation also causes a robust upregulation of an array of cell surface receptors. For this study, we quantified expression of CD69, the function of which is not fully understood, and CD25, the high-affinity subunit of the IL-2 receptor, both of which are early T cell activation markers. We also quantified CD44, which is constitutively expressed by memory T cells, but rapidly induced in naïve T cells following activation. The inducible costimulatory receptors, OX-40 and CD137, were also measured. In contrast to all the other markers that are upregulated following acitvation, CD62L, an adhesion molecule that directs T cells into lymph nodes, is rapidly downregulated following activation. In CD4 T cells, there was a decreased percentage of cells expressing the late activation markers, OX-40 and CD137, in the Bph/2 group (Fig. 7a). In contrast, there were no differences in the expression of the early activation markers CD69 and CD25 in CD4 T cells between the two strains. While we observed decreased expression of CD44 in resting CD4 T cells, we know from our previous analysis, that this is likely due to a decreased effector memory CD4 T cell population. In CD8 T cells, we observed a decreased percentage of activated T cells in Bph/2 mice, which was consistent across all the inducible markers (Fig. 7b).

**Figure 7.**
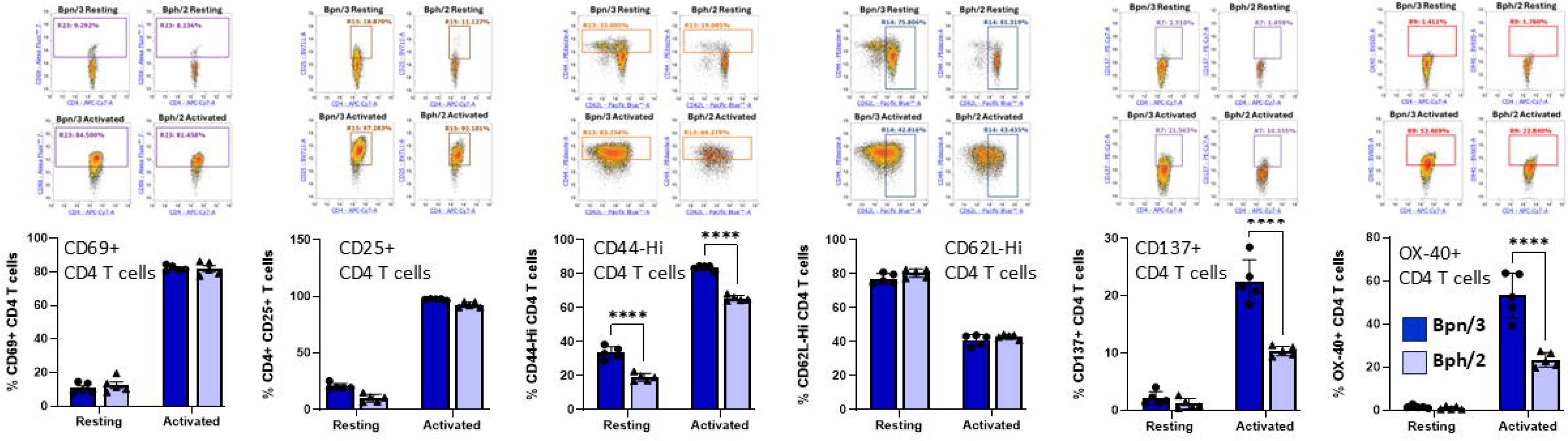
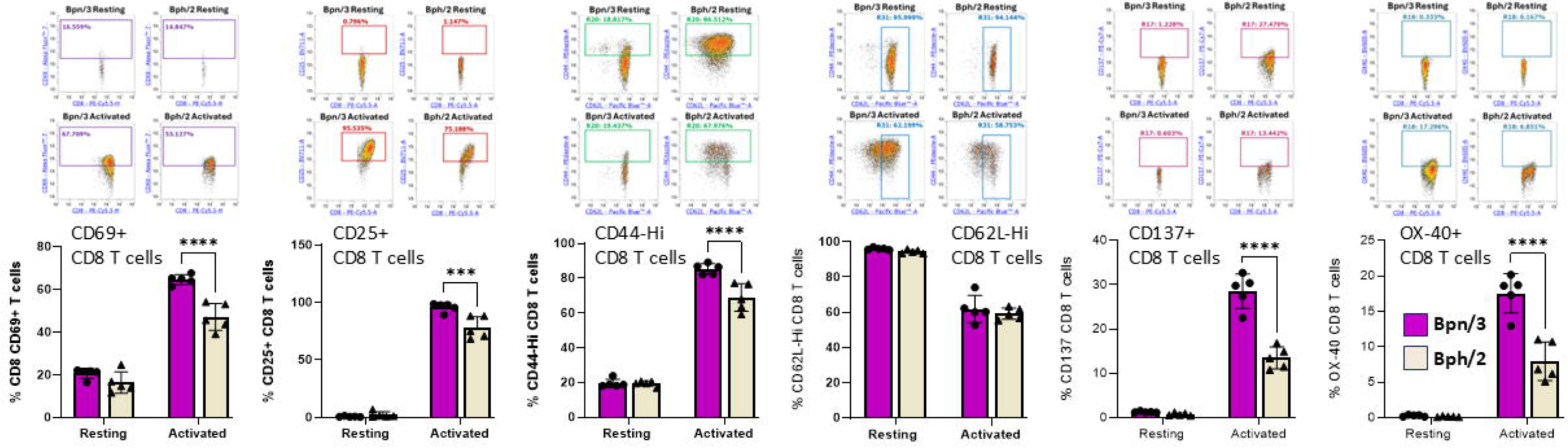
Diminished induction of T cell activation markers in CD8 T cells, and to a lesser extent, CD4 T cells from Bph/2 mice. Freshly isolated splenocytes were cultured in the presence or absence anti-CD3/anti-CD28 activating antibodies for 24h. The cells were then labeled with fluorochrome-conjugated antibodies (Table 1) and analyzed by flow cytometry. A) Decreased percentages of CD4^+^ T cells from BPH/2J mice were CD44^Hi^, CD137^+^, and OX-40^+^ 24-hours post-activation, whereas no differences were observed in percentages of CD25^+^, CD69^+^, and CD62L^Hi^. B) BPH/2J mice had decreased percentages of CD8^+^ T A) CD4 T cells and B) CD8 T cells were assessed for expression of the inducible cell surface markers: CD69, CD25, CD44, CD137 and OX-40. CD62L, which is rapidly downregulated after activation was also quantified. Above each plotted graph are representative dot plots. The data are presented as the mean ± SEM, n=5. * p<0.05 as determined by 2-way ANOVA followed by Dunnett’s post hoc analysis. ***p<0.001 and ****p<0.0001.

### Diminished early induction of most cytokines in activated splenocytes from Bph/2 mice

In correlation with upregulation of proliferation and activation markers, we all also assessed differences in cytokine production between the Bph/2 and Bpn/3 strains. Shortly after activation, T cells rapidly secrete an array of cytokines. CD4 T cells, in particular, differentiate into multiple functionally distinct subsets that are distinguished in part functionally by the cytokines produced. In assessing cytokines that are produced by T cells, we observed consistently decreased induction of cytokines in cells from Bph/2 mice 24 h after activation (Fig. 8). At this early time point, we did not observe differences in subsets associated with any one particular T cell lineage (Th1, Th2, Th17, etc.). The one exception to this was IL-9 which was not significantly decreased in the Bph/2 group and is associated with Th9 cells and a type 2 immune response. We also noted the presence of cytokines that are not typically produced in appreciable levels by T cells. These cytokines were likely produced by macrophages and other myeloid cells that were stimulated by cytokines and costimulatory factors produced by the activated T cells (Fig. 9). In addition, our data show upregulation of numerous chemokines, which are small cytokines that stimulate cell trafficking (Fig .10). While myeloid-derived cytokines were somewhat comparable between strains, we observed significantly lower secretion of IL-1α, IL-1β and VEGF in activated splenocytes from Bph/2 mice (Fig. 9). We also observed decreased levels of the MIP3α and RANTES chemokines in the Bph/2 group (Fig. 10). Because trends in cytokine production can change over time following activation, we also looked at T cell-derived cytokines at 120 h after activation. In general, the expression of many cytokines was largely equivalent between the two strains at this time point (Fig. 11). There were exceptions to this, however, including markedly increased IL-2, but decreased Th1 (IFNγ and TNFα) and Th17 (IL-17F, IL-21, IL-22, TNFβ) cytokines in the Bph/2 group. Overall, the data point to diminished induction of T cell cytokines in activated cells from Bph/2 mice at early time points after activation with a persistent decrease in Th1 and Th17 cytokines at late time points.

**Figure 8.**
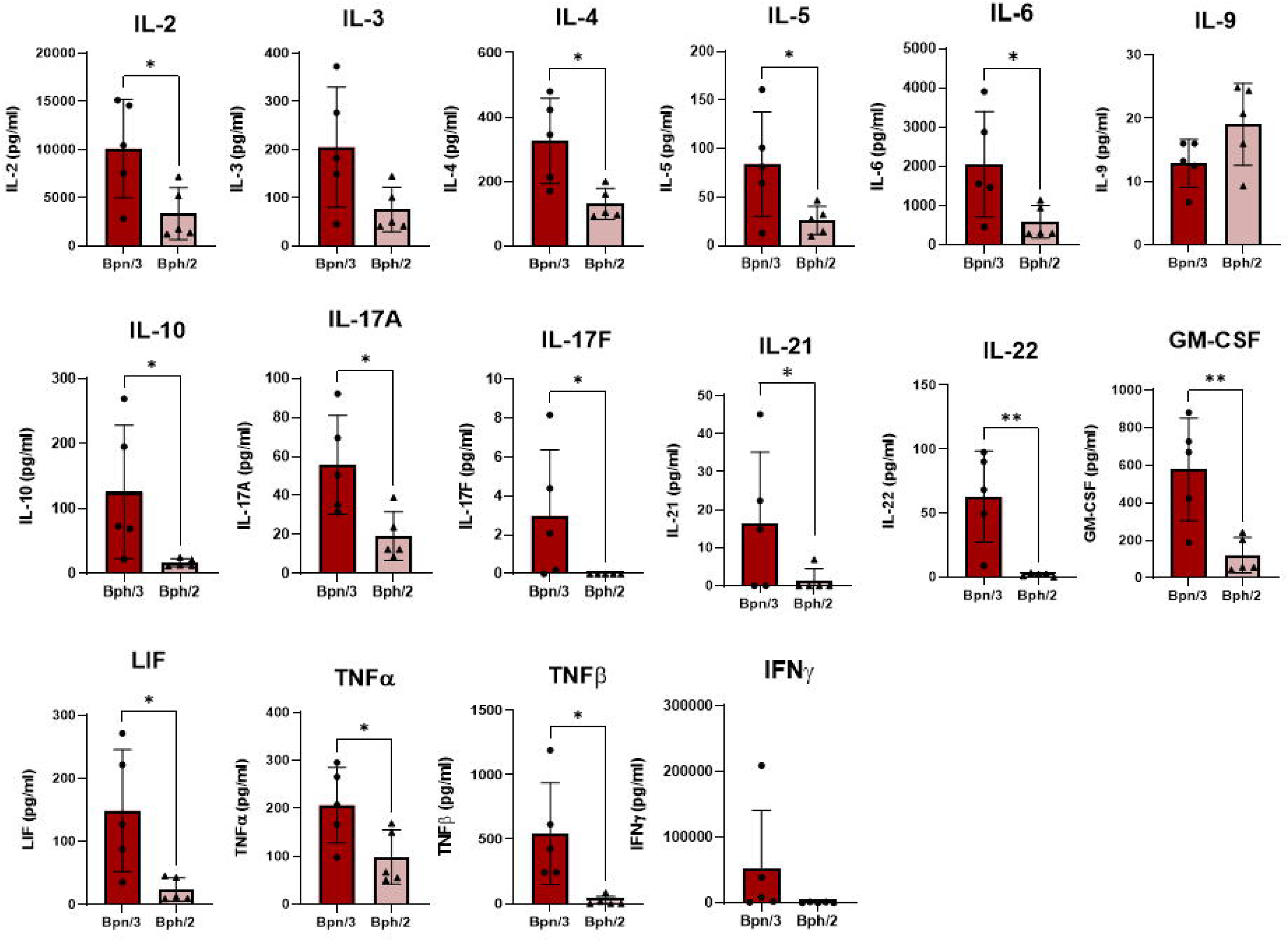
Reduced induction of T cell cytokines by splenic T cells from Bph/2 mice 24 h after activation. Freshly isolated splenocytes were cultured in the presence or absence anti-CD3/anti-CD28 activating antibodies. Cell supernatants were collected after 24 h for cytokine analysis by EVE Technologies (Calgary, Alberta, Canada). Data are presented as the mean ± SEM, n = 5. * p<0.05 and **p<0.01 as determined by 2-way ANOVA followed by Dunnett’s post hoc analysis.

**Figure 9.**
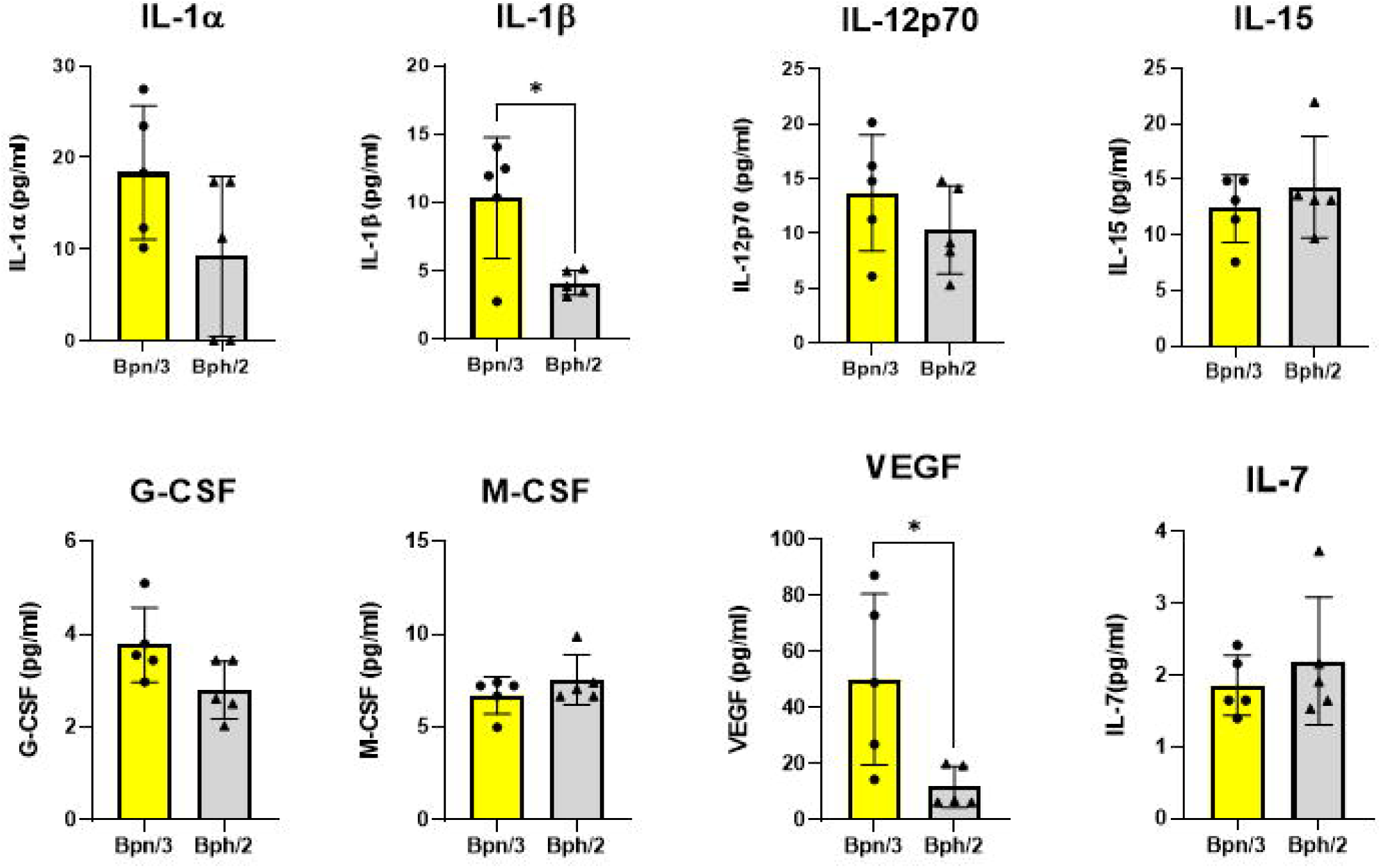
Decreased induction of IL-1β and VEGF by myeloid cells from Bph/2 mice 24 h after activation. Freshly isolated splenocytes were cultured in the presence or absence anti-CD3/anti-CD28 activating antibodies. Cell supernatants were collected after 24 h for cytokine analysis by EVE Technologies (Calgary, Alberta, Canada). Induction of cytokines was likely the result of indirect stimulation of myeloid cells by cytokines released by T cells. Data are presented as the mean ± SEM, n = 5. * p<0.05 as determined by 2-way ANOVA followed by Dunnett’s post hoc analysis.

**Figure 10.**
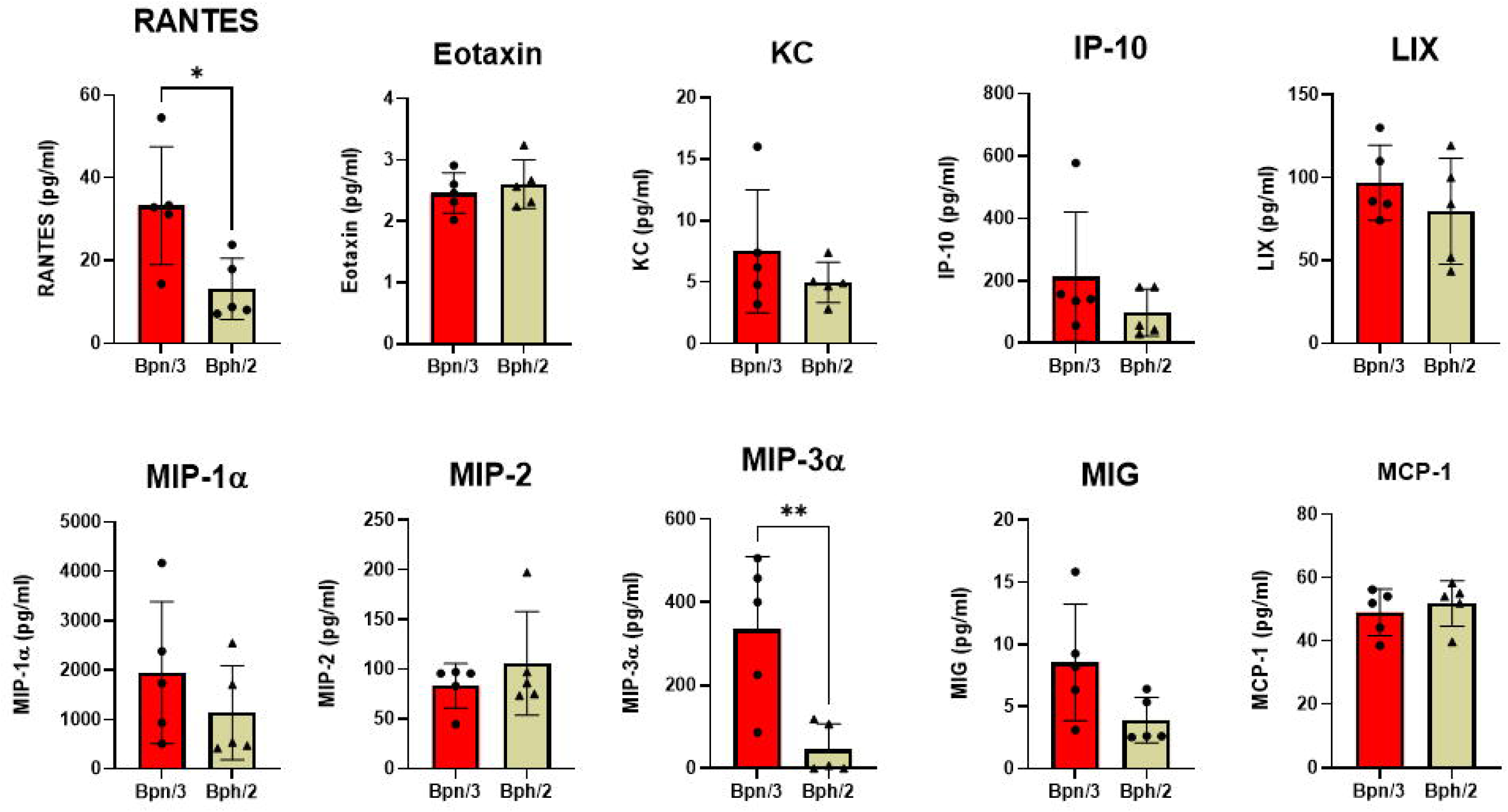
Reduced induction of the chemokines, RANTES and MIP3α, by splenic T cells from Bph/2 mice 24 h after activation. Freshly isolated splenocytes were cultured in the presence or absence anti-CD3/anti-CD28 activating antibodies. Cell supernatants were collected after 24 h for cytokine analysis by EVE Technologies (Calgary, Alberta, Canada). Data are presented as the mean ± SEM, n = 5. * p<0.05 and **p<0.01 as determined by 2-way ANOVA followed by Dunnett’s post hoc analysis.

**Figure 11.**
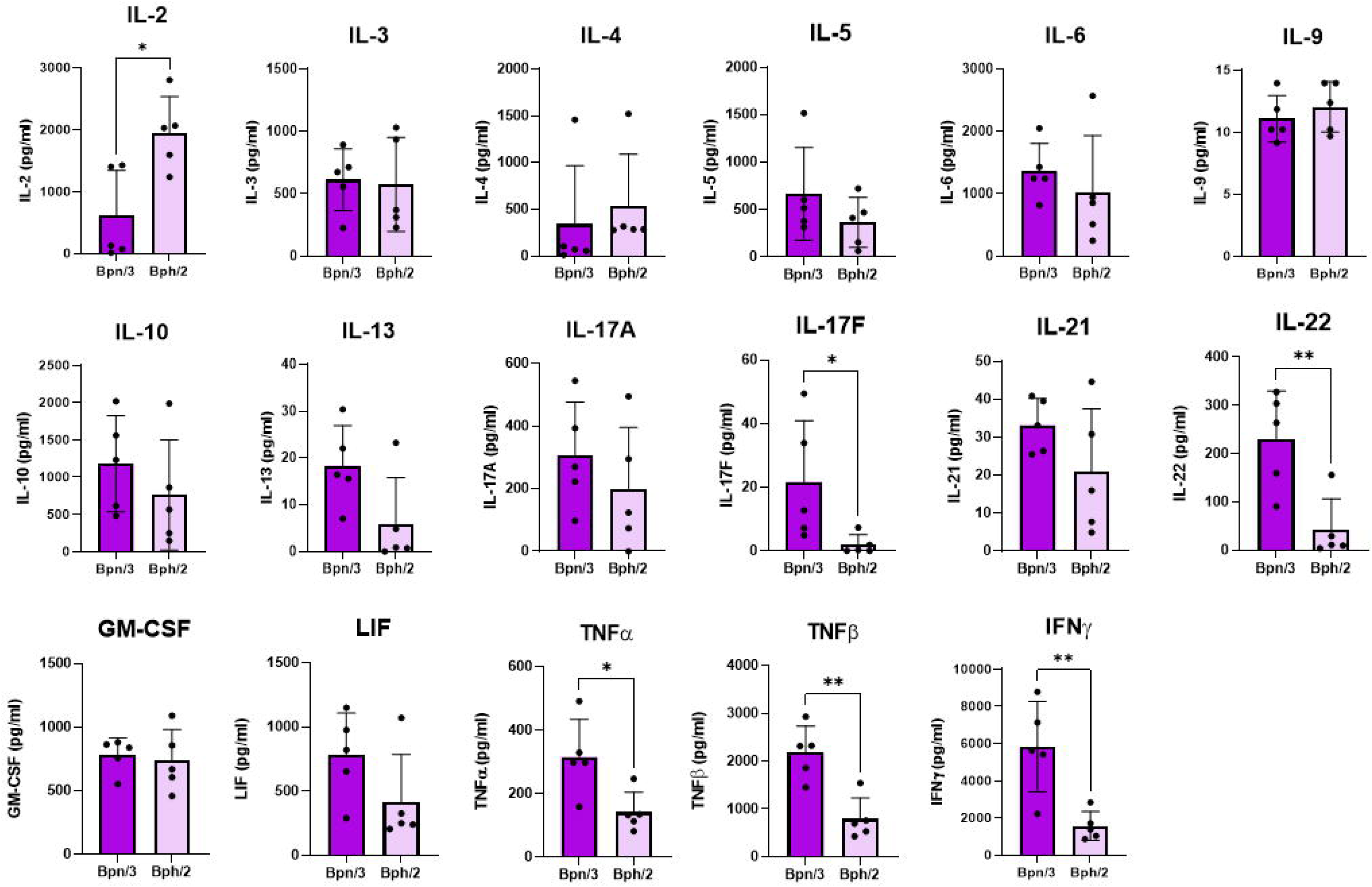
Decreased induction of T cell cytokines by splenic T cells from Bph/2 mice 120 h after activation. Freshly isolated splenocytes were cultured in the presence or absence anti-CD3/anti-CD28 activating antibodies. Cell supernatants were collected after 24 h for cytokine analysis by EVE Technologies (Calgary, Alberta, Canada). Data are presented as the mean ± SEM, n = 5. * p<0.05 and **p<0.01 as determined by 2-way ANOVA followed by Dunnett’s post hoc analysis.

## Discussion

The purpose of the present study was to compare the immune systems of the hypertensive Bph/2 mouse and normotensive Bpn/3 mice. Specifically, we compared immune cell composition in spleen and inguinal, brachial and mesenteric lymph nodes. We also assessed differences in lymphocyte response to polyclonal T cell activation. Overall, the immune cell composition was similar between the two strains with the exception of a markedly increased neutrophil population in the spleens of Bph/2 mice. While overall proportion of B cells was comparable between strains, there were differences in antibody expression in which there was a greater percentage of IgM+ B cells in Bph/2 mice that were either negative for, or had low expression of, IgD. We also noted a decreased percentage of effector memory cells within the CD4 T cell population and a decreased percentage of central memory cells within the CD8 T cell population. The diminished memory T cell populations correlated with decreased proliferation and a diminished cytokine response to polyclonal T cell activation. The decreased cytokine response was most pronounced at 24 h, but we also observed significantly diminished secretion of Th1 and Th17 cytokines at 120 h. Overall, the data indicate largely comparable immune cell composition between the two strains (with some exceptions), but notable functional differences in response to polyclonal T cell activation.

Dr. Gunther Schlager found significant differences in blood pressure between different strains of mice, which suggested a role for heritable genes in hypertension [9; 10; 11; 12; 13; 14; 15]. Accordingly, he originally generated the Bph/2 and the Bpn/3 mouse strains to identify the role of genetics in determining arterial blood pressure [1; 2; 16]. He found that blood pressures of the Bph/2 and Bpn/3 mice diverged as early as 6 weeks of age [1]. Similar to this report, we also observed significantly greater systolic, diastolic and mean arterial pressures in Bph/2 mice as compared to Bpn/3 mice. Dr. Schlager’s group also found an increase in heart rate. While our data show a trend toward an increase in heart rate in Bph/2 mice, the difference was not statistically significant. Notably, Jackson et al. has demonstrated that time of day plays a role in the magnitude of difference in blood pressure and heart rate between Bph/2 and Bpn/3 mice [17]. Of note, our measurements were taken during the day and Jackson et al. showed that the greatest differences in blood pressure and heart rate were observed at night. Nonetheless, we found significantly elevated blood pressure in Bph/2 mice, which is consistent with previous studies and use of Bph/2 as a hypertensive model.

The most pronounced difference in immune composition between the two mouse strains is in the percentage of splenic neutrophils, which was greater in Bph/2 mice. The biological significance of this is not clear, since neutrophils account for a fairly small fraction of immune cells in spleen. However, the increased percentage of neutrophils in spleen may be proportional to differences in the blood neutrophil population since one of the functions of the spleen is to filter blood. Indeed, an increased proportion of blood neutrophils has been reported in Bph/2 mice as compared to Bpn/3 mice [18]. It is also notable that these are data are consistent with a previous study that showed increased splenic neutrophils in diabetic and sham control Bph/2 mice as compared to Bpn/3 mice [19].

Although we did not observe differences in the percentage of B cells in any compartment within the Bpn/3 and Bph/2 strains, we did observe differences in the expression of antibodies that was specific to spleen. Antibody expression in B cells is dynamic and reflects in part different stages in B cell development. Whereas newly mature B cells in spleen and lymph nodes express high levels of IgD, IgD expression is downregulated following B cell activation while IgM expression is maintained [20; 21]. Some activated B cells may undergo class switching where both IgD and IgM expression are turned off and other antibody classes are expressed (IgG, IgA, etc. [22]. In addition to antibody production, B cells are also characterized by their anatomical location within lymphoid organs where the dominant population are the follicular B cells, which upon activation produce antibody in the follicles, and marginal zone B cells, which are innate-like B cells that provide quick responses to pathogens [23]. Our data showed a decrease in the percentage of IgM+ IgD^Hi^ B cells with a concurrent increase in percentage of IgM+IgD^Lo^ and IgM+ IgD-B cell populations. Thus, these data indicate a diminished proportion of naïve, unactivated B cells in the Bph/2 mice with a concurrent increased percentage of antigen-experienced B cells (IgM+ IgD-). The significance of the differences in the IgM+ IgD^Lo^ B cell population is not clear since there are several functionally-distinct subpopulations within that group, including T1 transitional cells (transitioning to naïve, mature phenotype), regulatory B cells and marginal zone B cells [24; 25; 26]. Of interest, we also observed an increase in the percentage of IgM+ IgD-Lo CD23-B cells (which includes the marginal zone B cell population) in the inguinal lymph nodes of Bph/2 mice [27]. In addition, we observed a decrease in IgM-IgD-B cells in the brachial lymph nodes of Bph/2 mice, which suggests decreased class switching to IgG and/or other antibody classes. However, this trend was not observed in the other compartments. We did not observe notable differences in the B cell activation markers CD69 and CD86 between the two strains, with the exception of in mesenteric lymph nodes, where CD86 was decreased in the Bph/2 mice. However, there was a fair amount of variability in the Bpn/3 group and that trend was not noted in the other compartments. Overall, the data suggest a greater percentage of B cells that have previously undergone activation in the spleens of Bph/2 mice.

Under homeostatic conditions (in the absence of exposure to a recent polyclonal activator), T cells can be defined as naïve (never activated) or as one of several memory populations [28]. While resident memory T cells are often found in tissues, such as lung and gut, effector memory and central memory T cells can be found in lymphoid organs [29; 30]. All memory T cells have a lower threshold for activation compared to naïve T cells. While effector memory T cells gain effector function quickly after activation, central memory T cells take more time to become effector T cells. Conversely, central memory T cells are more proliferative than effector memory T cells in response to activation. Both effector and central memory T cells are more proliferative and induce greater levels of cytokine expression than naïve T cells. Within the CD4 population, our data demonstrate a decreased percentage of effector memory T cells, which was consistent across compartments, but most pronounced in spleen and mesenteric lymph nodes. Conversely, in the CD8 T cell population, there was a consistent decrease in the percentage of central memory T cells across all compartments. The decreased percentage of memory T cells in Bph/2 mice is consistent with, and could be largely responsible for, the decrease in T cell proliferation and cytokine induction in response to polyclonal activation.

There are some limitations in this study. Because this was our first investigation into Bph/2 mice, we kept the study somewhat limited in scope. For example, we are not able to determine sex differences because we only included male mice in this study. However, we intend to follow up with a subsequent study that will include both sexes. Our intention was to conduct a comprehensive assessment of baseline immunology in multiple lymphoid compartments. While we generated considerable data in spleen and inguinal, brachial, mesenteric lymph nodes, it would be interesting to expand this to include immune cell composition in blood, bone marrow and other relevant tissues. To assess immune cell composition, we used multiple flow cytometry panels, which allowed us to assess broad immune cell composition as well as B cell and T cell subpopulations. Using this approach, we found notable differences in T cell memory populations between the Bph/2 and Bpn/3 strains. However, this leads to the question of why this difference occurs. The development of memory can be impacted by numerous factors, including the strength of the primary response in naïve T cells, T cell exhaustion and expression of transcription factors that regulate memory, for example [31; 32; 33]. None of these factors were addressed in the current study, but would be interesting to pursue in future studies. Likewise, the differences in B cell antibody production between the two strains could be due to differences in the threshold for B cell activation, which was not assessed in this study. Overall, the study pointed to a number of interesting immunological differences between the Bph/2 and Bpn/s mouse strains, but follow-up studies will be needed to understand why these differences occur.

The overall scientific question that this study aimed to address was whether Bph/2 and Bpn/3 mice share similar immune cell composition and similar T cell and B cell functionality. While the answer to the first question regarding immune cell composition is likely yes, the answer to the question of function is clearly no. There are notable functional differences in the lymphocyte populations. While the physiological significance of these differences is not yet clear—particularly with regard to the development of chronic diseases like hypertension, the data strongly suggest notable immune differences in these strains that could have long-term implications on immunity and inflammation.

## References

[1] G. Schlager, and J. Sides, Characterization of hypertensive and hypotensive inbred strains of mice. Lab Anim Sci 47 (1997) 288–92.

[2] G. Schlager, Selection for blood pressure levels in mice. Genetics 76 (1974) 537–49.

[3] T. Okuda, and A. Grollman, Passive transfer of autoimmune induced hypertension in the rat by lymph node cells. Tex Rep Biol Med 25 (1967) 257–64.

[4] F.N. White, and A. Grollman, Autoimmune Factors Associated with Infarction of the Kidney. Nephron 1 (1964) 93–102.

[5] T.J. Guzik, N.E. Hoch, K.A. Brown, L.A. McCann, A. Rahman, S. Dikalov, J. Goronzy, C. Weyand, and D.G. Harrison, Role of the T cell in the genesis of angiotensin II induced hypertension and vascular dysfunction. J Exp Med 204 (2007) 2449–60.

[6] D.L. Mattson, H. Lund, C. Guo, N. Rudemiller, A.M. Geurts, and H. Jacob, Genetic mutation of recombination activating gene 1 in Dahl salt-sensitive rats attenuates hypertension and renal damage. Am J Physiol Regul Integr Comp Physiol 304 (2013) R407–14.

[7] N. Rudemiller, H. Lund, H.J. Jacob, A.M. Geurts, D.L. Mattson, and P. PhysGen Knockout, CD247 modulates blood pressure by altering T-lymphocyte infiltration in the kidney. Hypertension 63 (2014) 559–64.

[8] R.A. Freeborn, A.P. Boss, L.M. Kaiser, E.M. Gardner, and C.E. Rockwell, Trivalent arsenic impairs the effector response of human CD4(+) and CD8(+) T cells to influenza A virus ex vivo. Food Chem Toxicol 165 (2022) 113122.

[9] R.S. Weibust, and G. Schlager, A genetic study of blood pressure, hematocrit and plasma cholesterol in aged mice. Life Sci 7 (1968) 1111–9.

[10] G. Schlager, and R.S. Weibust, Genetic control of blood pressure in mice. Genetics 55 (1967) 497–506.

[11] G. Schlager, Spontaneous hypertension in laboratory animals. A review of the genetic implications. J Hered 63 (1972) 35–8.

[12] G. Schlager, Genetic control of blood pressure by more than one pair of alleles. Proc Soc Exp Biol Med 136 (1971) 863–6.

[13] G. Schlager, Genetic and physiological studies of blood pressure in mice. I. Crosses between A-J, SWR-J, and their hybrids. Can J Genet Cytol 10 (1968) 853–64.

[14] G. Schlager, Systolic blood pressure in eight inbred strains of mice. Nature 212 (1966) 519–20.

[15] G. Schlager, Heritability of blood pressure in mice. J Hered 56 (1965) 278–84.

[16] G. Schlager, and C.S. Chao, The role of dominance and epistasis in the genetic control of blood pressure in rodent models of hypertension. Clin Exp Hypertens A 13 (1991) 947–53.

[17] K.L. Jackson, G.A. Head, C. Gueguen, E.R. Stevenson, K. Lim, and F.Z. Marques, Mechanisms Responsible for Genetic Hypertension in Schlager BPH/2 Mice. Front Physiol 10 (2019) 1311.

[18] M.R. Kandalgaonkar, B.S. Yeoh, B. Joe, N.W. Schmidt, M. Vijay-Kumar, and P. Saha, Hypertension Increases Susceptibility to Experimental Malaria in Mice. Function (Oxf) 5 (2024) zqae009.

[19] A. Sharma, J.S.Y. Choi, A.M.D. Watson, L. Li, T. Sonntag, M.K.S. Lee, A.J. Murphy, M. De Blasio, G.A. Head, R.H. Ritchie, and J.B. de Haan, Cardiovascular characterisation of a novel mouse model that combines hypertension and diabetes co-morbidities. Sci Rep 13 (2023) 8741.

[20] J.L. Preudhomme, Loss of surface IgD by human B lymphocytes during polyclonal activation. Eur J Immunol 7 (1977) 191–3.

[21] A. Bourgois, K. Kitajima, I.R. Hunter, and B.A. Askonas, Surface immunoglobulins of lipopolysaccharide-stimulated spleen cells. The behavior of IgM, IgD and IgG. Eur J Immunol 7 (1977) 151–3.

[22] J. Stavnezer, and C.E. Schrader, IgH chain class switch recombination: mechanism and regulation. J Immunol 193 (2014) 5370–8.

[23] T.W. LeBien, and T.F. Tedder, B lymphocytes: how they develop and function. Blood 112 (2008) 1570–80.

[24] F. Loder, B. Mutschler, R.J. Ray, C.J. Paige, P. Sideras, R. Torres, M.C. Lamers, and R. Carsetti, B cell development in the spleen takes place in discrete steps and is determined by the quality of B cell receptor-derived signals. J Exp Med 190 (1999) 75–89.

[25] A. Cerutti, M. Cols, and I. Puga, Marginal zone B cells: virtues of innate-like antibody-producing lymphocytes. Nat Rev Immunol 13 (2013) 118–32.

[26] S.D. Neu, and B.N. Dittel, Characterization of Definitive Regulatory B Cell Subsets by Cell Surface Phenotype, Function and Context. Front Immunol 12 (2021) 787464.

[27] T.J. Waldschmidt, F.G. Kroese, L.T. Tygrett, D.H. Conrad, and R.G. Lynch, The expression of B cell surface receptors. III. The murine low-affinity IgE Fc receptor is not expressed on Ly 1 or ’Ly 1-like’ B cells. Int Immunol 3 (1991) 305–15.

[28] C.R. Mackay, T-cell memory: the connection between function, phenotype and migration pathways. Immunol Today 12 (1991) 189–92.

[29] F. Sallusto, J. Geginat, and A. Lanzavecchia, Central memory and effector memory T cell subsets: function, generation, and maintenance. Annu Rev Immunol 22 (2004) 745–63.

[30] R.A. Clark, Resident memory T cells in human health and disease. Sci Transl Med 7 (2015) 269rv1.

[31] J.M. Angelosanto, S.D. Blackburn, A. Crawford, and E.J. Wherry, Progressive loss of memory T cell potential and commitment to exhaustion during chronic viral infection. J Virol 86 (2012) 8161–70.

[32] Y. Chen, R. Zander, A. Khatun, D.M. Schauder, and W. Cui, Transcriptional and Epigenetic Regulation of Effector and Memory CD8 T Cell Differentiation. Front Immunol 9 (2018) 2826.

[33] M.A. Daniels, and E. Teixeiro, TCR Signaling in T Cell Memory. Front Immunol 6 (2015) 617.

